# Machine learning determines stemness associated with simple and basal-like canine mammary carcinomas

**DOI:** 10.1101/2023.10.09.561566

**Authors:** Pedro Luiz P. Xavier, Maycon Marção, Renan L. S. Simões, Maria Eduarda G. Job, Ricardo de Francisco Strefezzi, Heidge Fukumasu, Tathiane M. Malta

## Abstract

Simple and complex carcinomas are the most common type of malignant Canine Mammary Tumors (CMTs), with simple carcinomas exhibiting aggressive behavior and poorer prognostic. Stemness is an ability associated with cancer initiation and progression, malignancy, and therapeutic resistance, but is still few elucidated in different canine mammary cancer subtypes. Here, we first validated, using CMT samples, a previously published canine one-class logistic regression machine learning algorithm (OCLR) to predict stemness (mRNAsi) in canine cancer cells. Then, we observed that simple carcinomas exhibit higher stemness than other histological subtypes and confirmed that stemness is higher and associated with basal-like tumors and with NMF2 tumor-specific metagene signature. Furthermore, using correlation analysis, we suggested two promise stemness-associated targets in CMTs, *POLA2* and *APEX1*, especially in simple canine mammary tumors. Thus, our work elucidates stemness as a potential mechanism behind the aggressiveness and development of simple canine mammary tumors, describing novel pieces of evidence of a promising strategy to target canine mammary carcinomas, especially the simple subtype.

## Introduction

Canine mammary tumors (CMTs) are the most common type of cancer in female dogs, and have been widely studied as a comparative model for breast cancer in women (Abdelmegeed and Mohammed 2018; Gray et al. 2020). Both share similar biological and clinical patterns, including a high incidence, hormonal influence, and histopathological profiles. Additionally, some similarities in molecular and genetic makeup can be observed, such as copy number variation in important breast cancer-associated genes such as *PTEN, MYC, BRCA1*, and *BRCA2* (Rivera et al. 2009; Borge et al. 2015). The histological features of mammary tumors in dogs and humans can be different, with benign tumors being more prevalent in CMTs (Salas et al. 2015). Furthermore, CMTs often exhibit mesenchymal tumors, such as fibrosarcomas and carcinosarcomas, and the proliferation of myoepithelial cells in complex adenomas and carcinomas, extremely rare in human breast cancers (Cassali et al. 2007; Goldschmidt et al. 2011).

Simple carcinomas are composed of only one cell population, resembling either luminal epithelial or myoepithelial cells, whereas complex carcinomas resemble two different cell populations: malignant epithelial luminal cells and myoepithelial cells. Dogs with simple carcinomas have poorer prognoses than those with other subtypes (Goldschmidt et al. 2011; Cassali et al. 2011; Karayannopoulou et al. 2005; Rasotto et al. 2017). However, the mechanisms underlying the association between simple CMTs and poorer prognosis remain unclear, and few studies have investigated this. In recent years, omics-based approaches have helped advance knowledge regarding the genetic and molecular landscape of these types of CMT. Liu et al. (Liu et al. 2014), using whole-exome sequencing (WES) and transcriptomics (RNA-Seq), identified and suggested that simple CMT probably arises from genomic aberrations, whereas complex CMT originates from epigenomic alterations. More recently, well-designed studies have traced molecular oncogenic signatures in CMT, identifying relevant mechanisms that would lead to a better understanding of the underlying molecular pathogenesis of CMT and comparing it with human breast cancer (Kim et al. 2020; Graim et al. 2021).

Stemness is the ability of stem cells (SC) to self-renew and differentiate, ensuring the maintenance of at least one new undifferentiated SC, but also to differentiate into mature cells of different adult organs (Lathia and Liu 2017). It is also a phenotype associated with the cancer stem cell (CSC) concept and is consequently interconnected with intratumoral heterogeneity, initiation, progression, multidrug resistance, and metastasis (Ayob and Ramasamy 2018). Some studies have elucidated the role of stemness in CMT therapies and histogenesis in dogs (Michishita, 2020; Marção *et al*., 2022). In addition, several studies have isolated cancer stem cells in different types of canine cancers, including osteosarcoma (Wilson et al. 2008), glioblastoma (Stoica et al. 2009), prostate cancer (Moulay *et al*., 2013), hepatocellular carcinoma (Michishita et al. 2014), hemangiosarcoma (Aoshima et al. 2018), and CMT (Michishita et al. 2012; Cocola et al. 2009). However, the influence of stemness on different CMT histological and molecular subtypes, such as simple and complex or basal-like and non-basal-like carcinomas, is still a largely unexplored field, and innovative findings could be exploited for the understanding of CMT histogenesis and the development of potential therapeutic approaches.

Here, we applied a one-class logistic regression (OCLR) machine-learning algorithm (Malta et al. 2018a; Marção et al. 2022) to extract transcriptomic feature sets from a public database and measure stemness in different subtypes of CMT. Our study observed that CMT samples had higher stemness than normal samples, validating our model. Furthermore, simple CMTs exhibited higher stemness than the other histological subtypes. We also confirmed that stemness is higher in basal-like samples and in a “DNA-repair” tumor-specific metagene signature previously determined (Kim et al. 2020). Finally, we correlated and suggested two potential targets, *POLA2* and *APEX1*, and target-specific compounds that may inhibit stemness in CMT, especially in the most aggressive subtypes.

## Results

### Stemness index is higher in malignant CMT in comparison with matched normal samples

In our previous study, we created and validated a canine prediction model for stemness using public data composed of canine non-tumor somatic and pluripotent stem cells. In addition, we applied this stemness model to different canine mammary tumor cell lines established in our laboratory (Marção et al. 2022). In the present study, we applied the canine stemness index algorithm to a public RNA-seq dataset comprising CMT and pairwise normal tissues. The dataset was from a cohort containing 157 cases of benign and malignant CMT, and 64 cases of matched normal CMT tissue (Kim et al. 2020). As a first step, we rebuilt the prediction model using the overlapping genes between the original stemness prediction model for human (Malta et al. 2018b) and the canine dataset. The efficiency of conversion of canine to human gene ID was approximately 45% (16,405/ 35,000 genes). By comparing the tumor and matched normal sequencing data, we observed that malignant tumors in general presented a higher stemness index (mRNAsi) than normal samples, but not in comparison with benign tumors (**Figure 1A-B**, *p* < 0.0001). Thus, these results further validated our canine prediction model of stemness in tissue samples and indicated that stemness is a relevant phenotype in malignant CMT.

**Figure 1.**
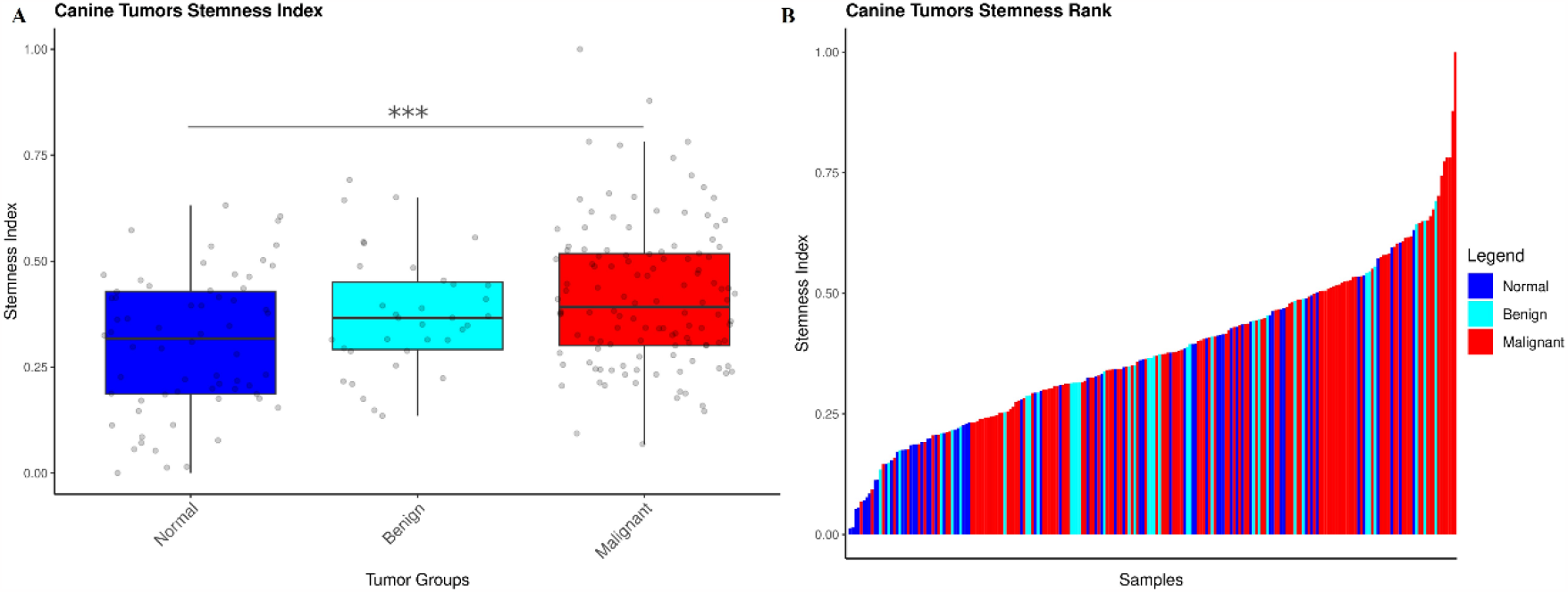
Stemness is higher in malignant CMTs in comparison with pairwise normal samples. **(A)** mRNAsi of normal samples and malignant and benign CMTs visualized in a boxplot and stratified by a median. Malignant CMTs exhibited the highest mRNAsi in comparison to pairwise normal samples. ^***^ = p < 0.001 (one-way ANOVA). **(B)** Rank plot corresponding to the mRNAsi predicted to each sample comparing malignant and benign CMTs and normal samples.

### Simple carcinomas exhibit higher stemness than other histologic subtypes

Clinically, the presence of myoepithelial cells in complex CMTs is associated with less aggressive biological behavior and better prognosis than simple CMTs (Karayannopoulou et al. 2005; Goldschmidt et al. 2011). To determine whether stemness is related to these clinical phenotypes, we applied mRNAsi in both histological subtypes and others, including carcinoma in benign mixed tumors, carcinosarcoma, fibrosarcoma, osteosarcoma, and spindle cell carcinoma. Interestingly, we observed that simple CMTs presented higher stemness than complex and other subtypes of CMT (**Figure 2A)**, with both the solid and tubulopapillary simple CMTs type exhibiting higher mRNAsi (**Figure 2B**). In addition, using correspondence analysis (CA), we demonstrated that high stemness was strongly associated with simple carcinomas while low stemness is associated with complex carcinomas **(Figures 2C and 2D**). Finally, pairwise analysis showed that simple carcinomas were the only subtype in which tumor samples exhibited significantly higher mRNAsi levels compared to matched normal tissues (**Figures 2E and 2F**).

**Figure 2.**
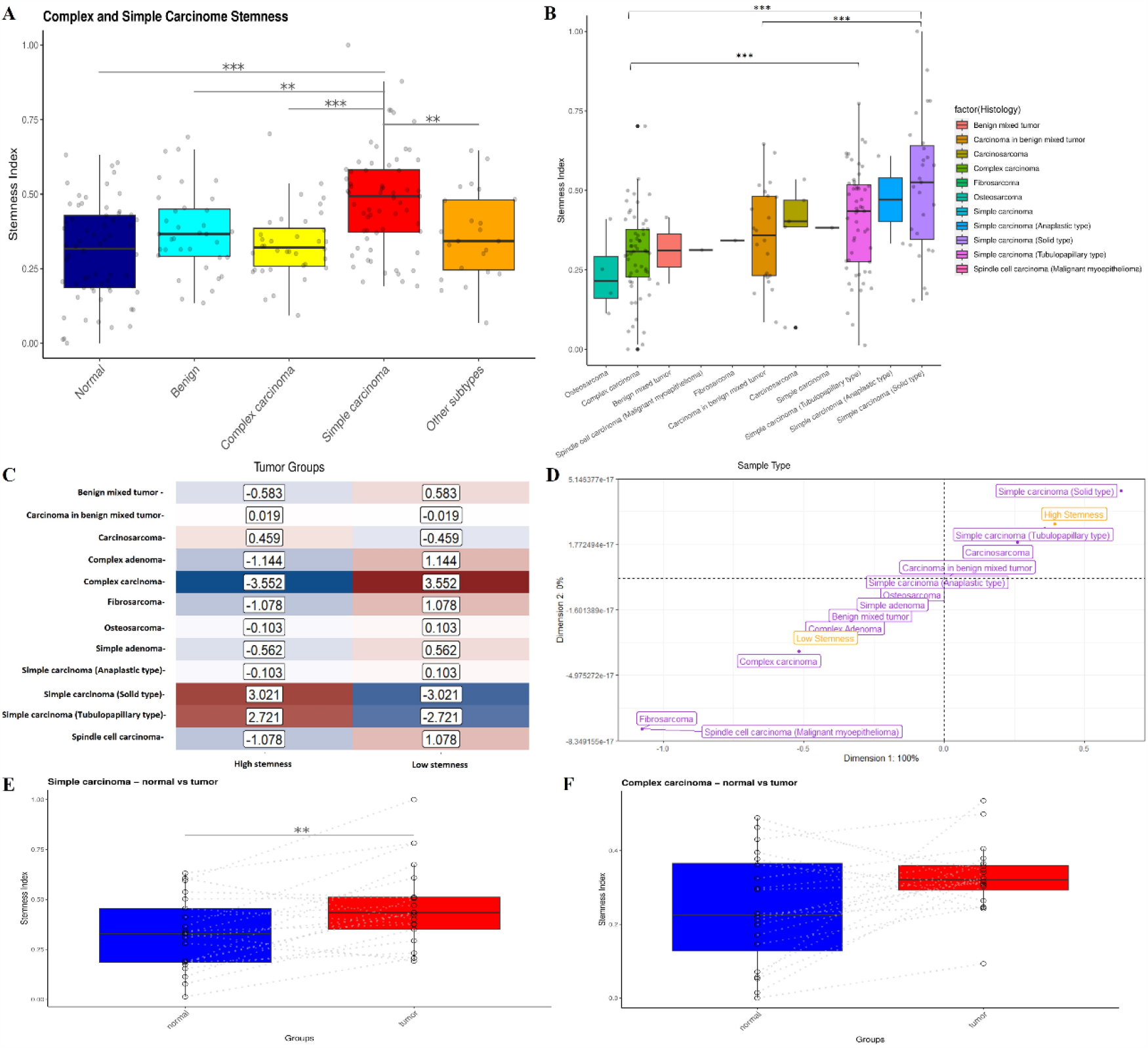
Stemness is higher in simple CMTs in comparison with complex and other CMT subtypes. (**A**) mRNAsi of CMT subtypes visualized in a boxplot and stratified by a median. Simple CMTs exhibited the highest mRNAsi in comparison to normal, benign, complex, and other subtypes of CMT. ^***^ = p < 0.001; ^**^ = p < 0.01 (one-way ANOVA). (**B**) mRNAsi is higher in solid simple CMTs in comparison to tubulopapillary simple CMT and other subtypes (one-way ANOVA). (**C**) Heatmap exhibiting the values of the adjusted standardized residuals. Categories of variables with values higher than 1.96 are associated. We could observe a strong association between low stemness and complex CMT (4.447) and between high stemness and simple CMT (4.914). (**D**) A CA two-dimensional perceptual map demonstrating the association between the categories of each categorical variable. Categories in yellow are stemness levels. Categories in purple are CMT Subtypes. Categories that are closely clustered are strongly associated with. (**E**) The pairwise analysis demonstrated that tumor simple CMT exhibits higher stemness than their matched normal samples. ^***^ = p < 0.001; ^**^ = p < 0.01 (T-test). (**F**) The pairwise analysis demonstrated that tumor complex CMT and their matched normal samples exhibit no statistical difference regarding stemness (T-test).

Using multiple correspondence analysis (MCA), we also evaluated whether simple and complex carcinomas were associated with clinical prognostic factors available in the database. Simple carcinomas were associated with tumor classification grade 3, negative status for estrogen-receptor expression, PAM50 classification of basal-like tumors, lymphatic invasion, and high stemness measured with mRNAsi. In contrast, complex carcinoma samples were associated with a positive status for estrogen receptor expression, grade 1, PAM50 classification of non-basal-like tumors, lymphatic invasion, and low stemness measured with mRNAsi (**Figure 3)**. Thus, these findings suggest that simple carcinomas are associated with poor clinical prognostic factors and that the stemness phenotype may contribute to their malignancy.

**Figure 3.**
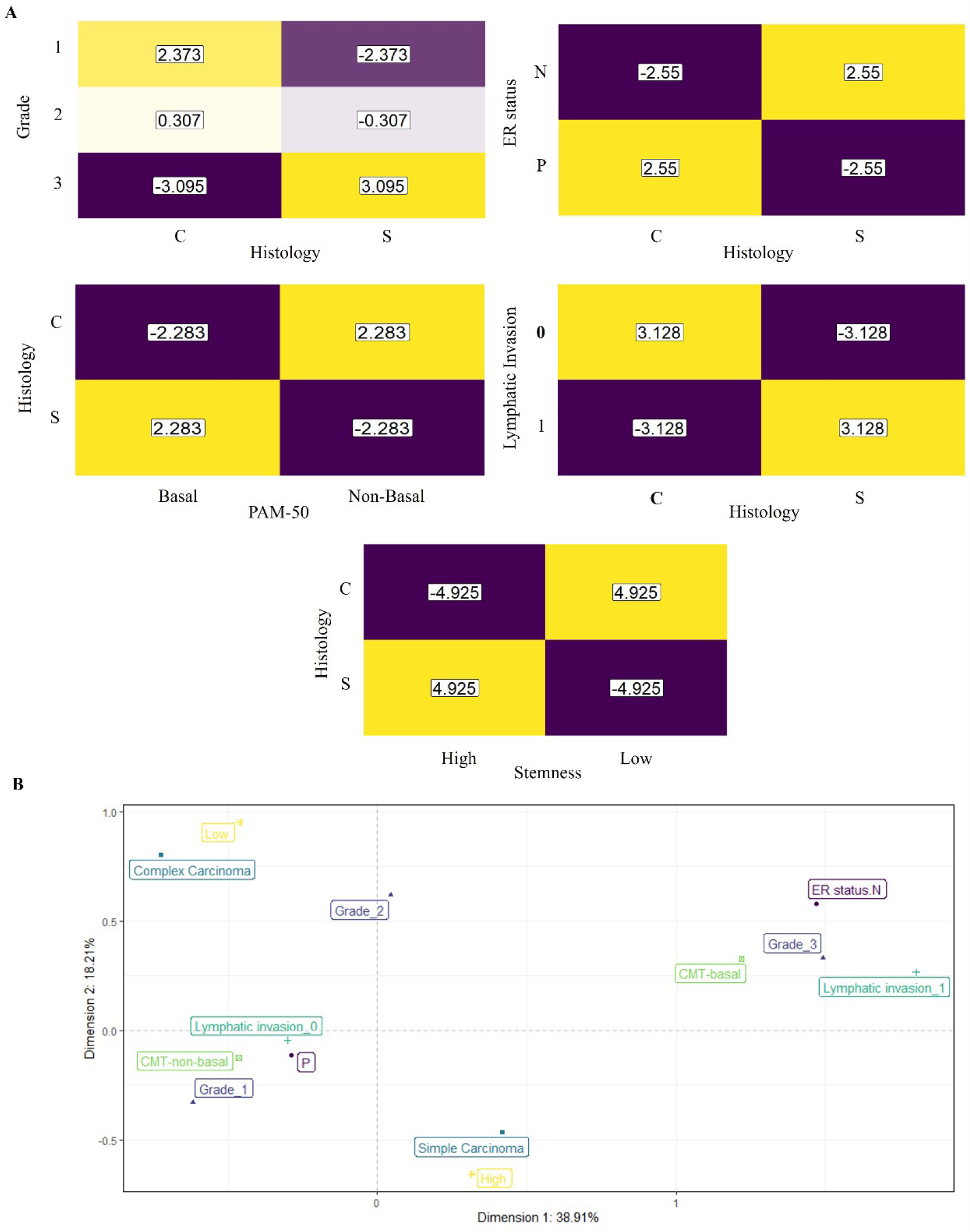
Simple CMTs are associated with poor prognostic factors. (**A**) Heatmap exhibiting the values of the adjusted standardized residuals. Categories of variables with values higher than 1.96 are associated. We could observe a strong association between simple CMT with poor prognostic variables such as tumor classification grade 3 (3.095), negative status for estrogen-receptor expression (2.55), PAM50 classification of basal-like tumors (2.283), lymphatic invasion (3.128), and high stemness (4.925). (**B**) A MCA two-dimensional perceptual map demonstrating the association between the categories of each categorical variable. Categories in yellow are stemness levels. Categories in blue are tumor classification grades. Categories in purple are the status for estrogen-receptor expression. Categories in light green are the PAM50 classification. Categories in green are the status of lymphatic invasion. Categories in dark green are CMT subtypes. Categories that are closely clustered are strongly associated with.

### High stemness index is associated with CMT-Basal and NMF2 molecular subtypes

We also investigated whether the stemness phenotype is associated with two different gene expression-based CMT subtypes according to PAM50 and NMF classifications. First, we measured the mRNAsi of CMT tumors classified as CMT-basal or CMT-non-basal, according to Kim *et al*. (Kim *et al*., 2020). We found that CMT-basal tumors presented higher stemness than CMT-non-basal tumors and, interestingly, only malign samples presented this difference, while no significant differences were found in pairwise normal samples and benign samples (**Figure 4A**). To further confirm this, we applied simple correspondence analysis and observed that high stemness was strongly associated with CMT-basal in malignant tumors (**Figure 4B**).

**Figure 4.**
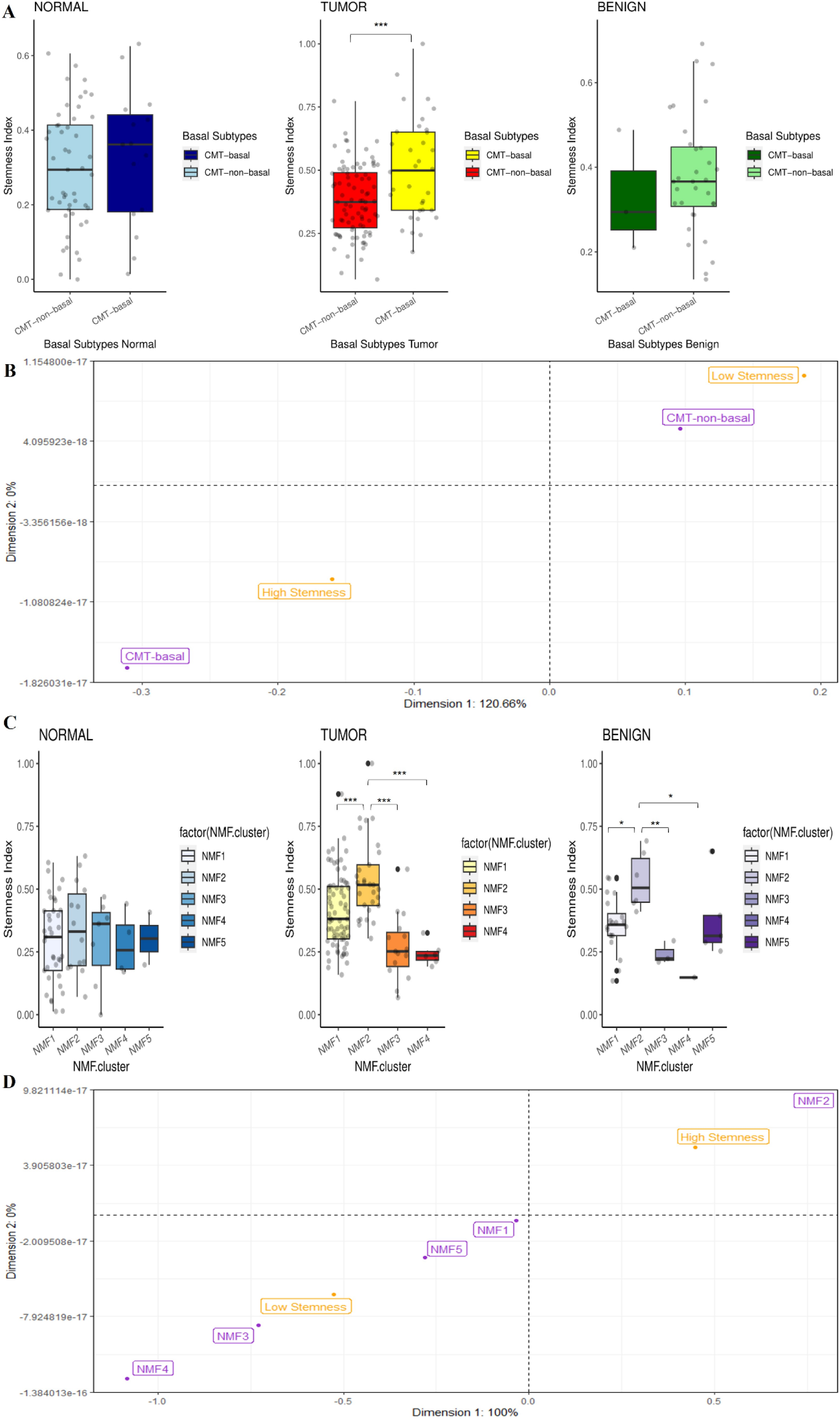
Stemness is higher in basal-like and NMF2 molecular subtypes. (**A**) mRNAsi visualized in a boxplot and stratified by a median. Basal-like CMTs exhibited the highest mRNAsi in comparison to non-basal subtypes of CMT. ^***^ = p < 0.001 (T-test (**B**) A CA two-dimensional perceptual map demonstrating the association between the categories of each categorical variable. Categories in yellow are stemness levels. Categories in purple are PAM50 Subtypes. Categories that are closely clustered are strongly associated with. (**C**). NMF2 CMTs exhibited the highest mRNAsi in comparison to NMF1, NMF3, and NMF4 subtypes both in malignant and benign tumors. ^***^ = p < 0.001; ^**^ = p < 0.01; ^*^ = p < 0.05 (one-way ANOVA) (**D**) A CA two-dimensional perceptual map demonstrating the association between the categories of each categorical variable. Categories in yellow are stemness levels. Categories in purple are NMF molecular Subtypes. Categories that are closely clustered are strongly associated with.

Kim *et al*. identified three gene expression-based CMT subtypes associated with molecular signatures using gene set enrichment analysis. Based on enriched molecular terms, the authors annotated three tumor-specific metagene signatures:1) “mitosis” (NMF1), “DNA repair” (NMF2), and “epithelial-to-mesenchymal transition” (EMT) (NMF3). Here, we assessed whether these metagene signatures were associated with the stemness index. We observed that NMF2 was the metagene signature with the highest stemness, both in malign and benign tumors (**Figure 4C**). We also confirmed these findings using correspondence analysis for malignant tumors (**Figure 4D**). Interestingly, 51% of samples classified as NMF2 were simple CMTs (18/35). Although NMF3 CMTs were shown to be a claudin-low subtype, known to be enriched with tumor-initiating cells or stem cells (Kim et al. 2020), our analysis demonstrated that the metagene signature NMF2 exhibits a higher potential of stemness.

### Correlation analysis identifies potential targets and inhibitors targeting stemness signature

We employed correlation analysis to identify correlations between genes, phenotypes, and compounds/inhibitors and to determine potential markers and targets in CMT, especially in high-stemness simple carcinomas. We found that 50 genes correlated with higher stemness, and 50 genes correlated with lower stemness (**Figure 5A**). Genes correlated with higher stemness were enriched for biological processes, such as DNA replication, cell division, cell cycle, and centromere complex assembly other terms associated with cell cycle (**Supplemental Table S1**). Genes correlated with lower stemness were enriched for biological processes, including the regulation of angiogenesis, cell migration, kinase activity, and cell adhesion (**Supplemental Table S2**). Two stemness-associated genes, *POLA2* and *APEX1*, are druggable targets for Dacarbazine and Lucanthone small inhibitors, respectively. Dacarbazine is an antineoplastic agent used to treat malignant melanoma and Hodgkin disease in humans (Chapman et al. 1999; Tarhini and Garwala 2006; Torka et al. 2022; Abramson et al. 2023). Lucanthone inhibits post-radiation DNA repair in tumor cells, especially in brain cancer. Furthermore, Lucanthone has been shown to inhibit stemness in glioma (Radin et al. 2022). An anticancer small molecule, APX-3330, also reduced colon cancer stem cell (CCSCs) growth in vitro. In addition, in xenograft mice injected subcutaneously with CCSCs, intratumoral administration of APX-3330 increased the tumor response to 5-Fluorouracil delivered intraperitoneally (Lou et al. 2014).

**Figure 5.**
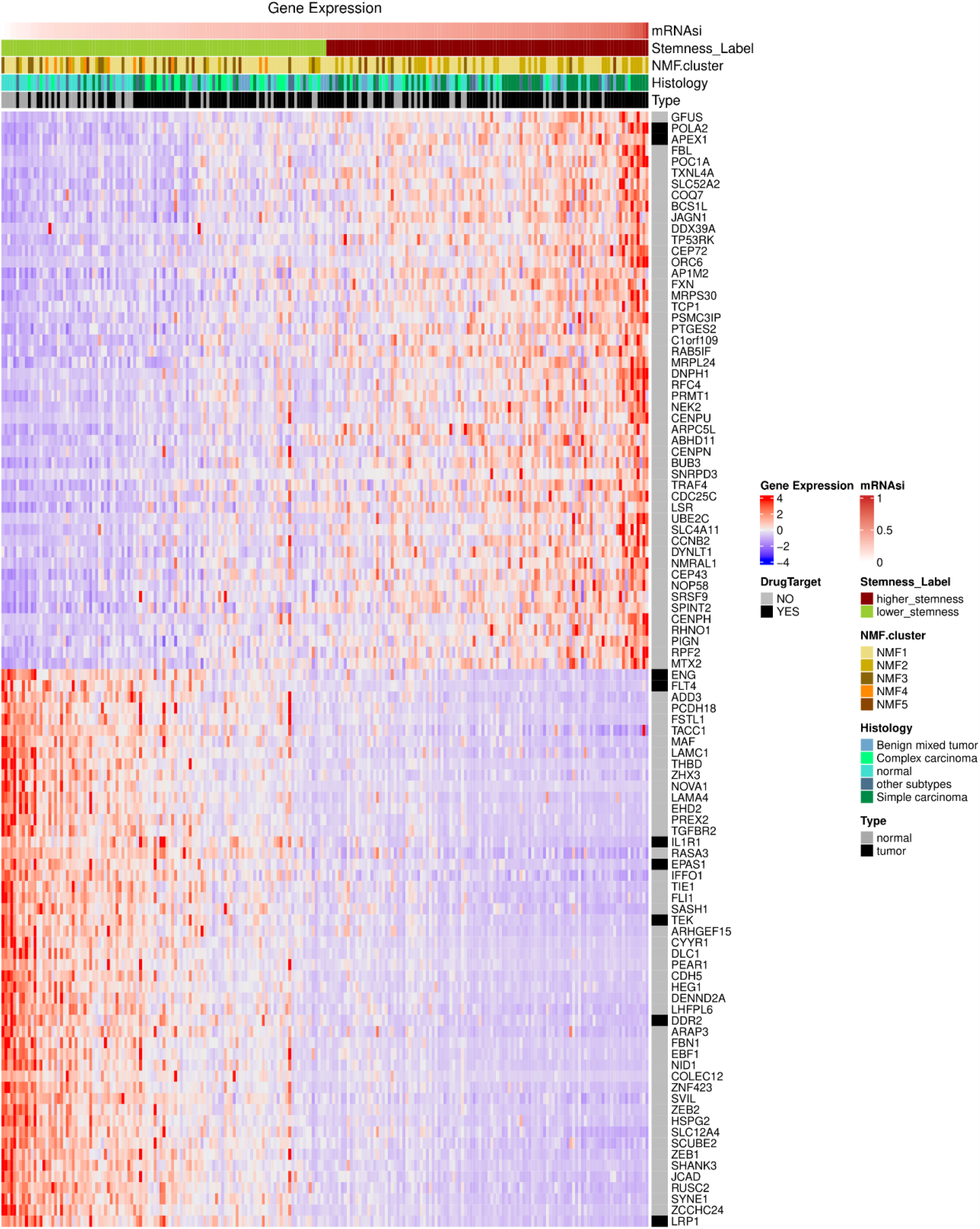
Correlation of Stemness, NMF cluster, Histology and sample type with target genes expression. Heatmap of canine mammary carcinoma gene expression data. Samples were sorted in ascending order (from left to right) according to mRNAsi. Above is the annotation of other parameters such as the categorical Stemness Index (high or low), NMF cluster designated for that sample, histological subtype of the tumor and indication of tumor or normal sample. On the right are listed the 100 genes (50 with the highest positive correlation and 50 with the highest negative correlation) related to the mRNAsi and which are anticancer drug targets according to the Open Targets platform.

We also demonstrated that *POLA2* is highly expressed in simple CMTs in comparison with normal samples (**Figure 6A**) and in the high-stemness NMF2 metagene signature (**Figure 6B)**. Furthermore, *APEX1* was highly expressed in simple carcinomas and other malignant subtypes in comparison with normal samples (**Figure 6C**) and in the NMF2 metagene signature (**Figure 6D**) demonstrating that both genes may be potential therapeutic targets in simple carcinomas and in high-stemness NMF2 samples.

**Figure 6.**
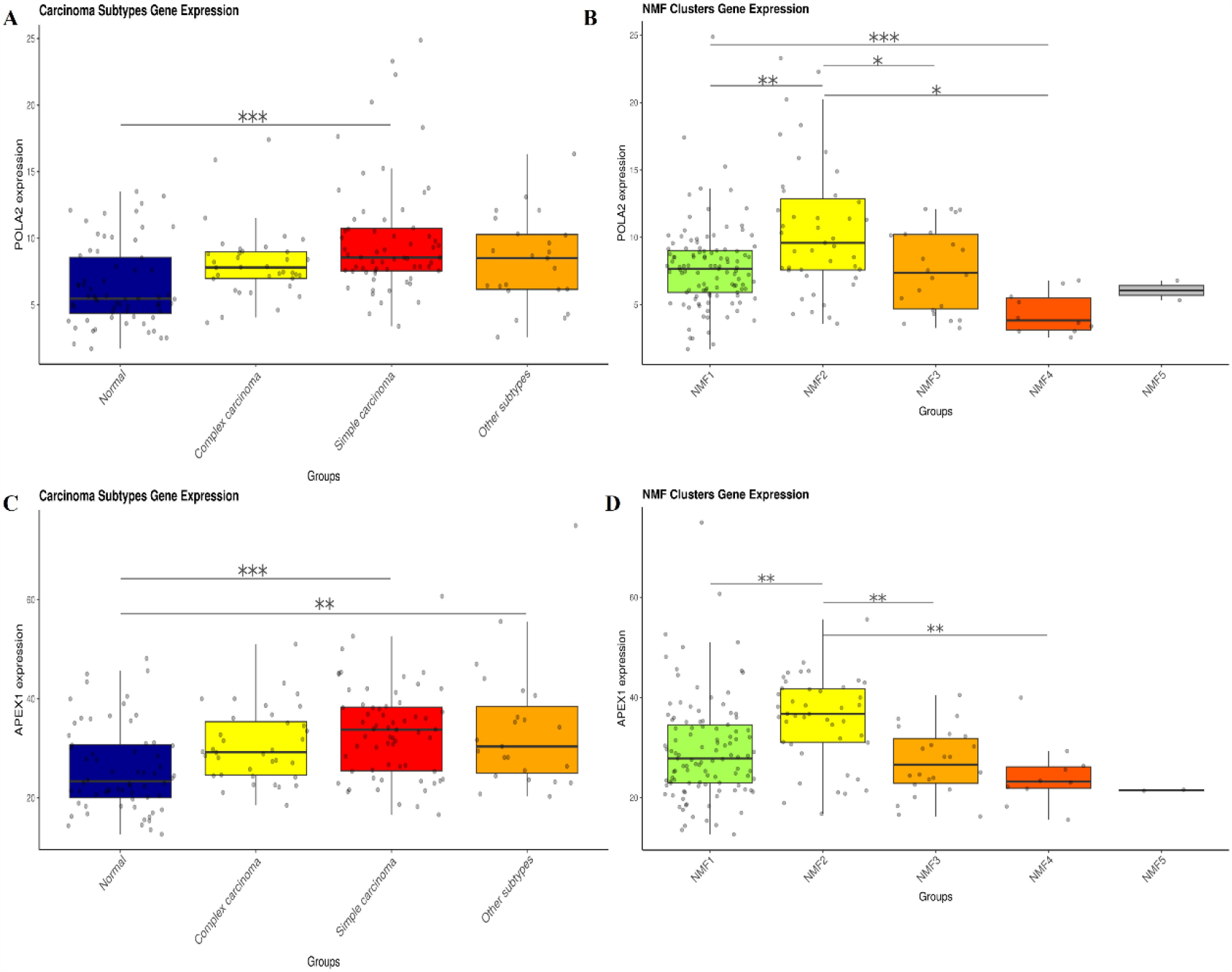
*POLA2* and *APEX1* expression are higher in simple CMTs and NMF2 metagene signature. (**A**) mRNAsi visualized in a boxplot and stratified by a median. Simple CMTs showed higher *POLA2* gene expression in comparison with normal samples. **(B)** mRNAsi visualized in a boxplot and stratified by a median. *POLA2* expression was higher in NMF2 metagene molecular subtype. (**C**) mRNAsi visualized in a boxplot and stratified by a median. *APEX1* was higher expressed in simple CMTs and other malignant subtypes in comparison with normal samples. **(D)** *APEX1* expression was higher in NMF2 molecular subtype. ^***^ = p < 0.001; ^**^ = p < 0.001; ^*^ = p < 0.05 (one-way ANOVA)

## Discussion

Stemness is a phenotype associated with cancer stem cells and has been implicated in cancer progression, malignancy, and tumor resistance. However, its roles in canine mammary carcinomas and their different subtypes, such as simple and complex, are poorly defined, partly owing to the lack of models to evaluate and measure stemness. Here, using a one-class logistic regression (OCLR) machine-learning algorithm to quantify cancer stemness and generate a stemness index (mRNAsi) in canine cancer (Malta *et al*., 2018; Marção *et al*., 2022), we found that stemness increased in malignant CMT samples in comparison with normal samples. In addition, we observed that stemness was associated with simple CMT and not with complex CMT, in addition to being associated with basal-like CMTs and the DNA repair metagene signature (Kim et al. 2020).

Despite the isolation of cancer stem cells from canine tumors in many studies (Wilson et al. 2008; Stoica et al. 2009; Moulay et al. 2013; Michishita et al. 2014; Aoshima et al. 2018; Michishita et al. 2012; Cocola et al. 2009), further investigation is needed to determine the key role of stemness in canine tumors. Advancements in omics-based technologies and databases have enabled comprehensive phenotypic and molecular studies of canine species (Kim et al. 2020; Graim et al. 2021; Liu et al. 2014). Using these public data, we could access and measure stemness in CMT and normal samples, suggesting interesting influences of stemness, especially in simple CMT. Simple CMT exhibit proliferation of only epithelial cells and have a worse prognosis than complex CMTs, but there is little evidence explaining this fact. Simple CMTs can faithfully recapitulate breast cancer in women and are closely clustered with basal-like human breast cancer in the PAM50 classification, suggesting a reason for their malignancy (Liu et al. 2014; Bergholtz et al. 2022; Watson et al. 2023). Our results also provide evidence to explain the malignancy of simple CMTs based on high stemness and, consequently, enrichment of cancer stem cells in these tumors. To date, no studies have demonstrated that canine simple CMT is enriched for cancer stem cells in comparison with other subtypes. However, several studies on human breast cancer have demonstrated an association between basal-like tumors, stemness, and cancer stem cells (Kim et al. 2015; Honeth et al. 2008; Zhou et al. 2019), contributing to the poor prognosis associated with this subtype (Yu et al. 2013), although the presence of cancer stem cells is not confined to basal-like tumors (Kim et al. 2012). Thus, our findings provide the first evidence that stemness may play key roles in simple CMTs; however, further well-designed studies involving the isolation and enrichment of cancer stem cells and validation of stemness markers in simple CMTs are needed to test this hypothesis.

Basal-like CMTs were associated with higher levels of stemness. To the best of our knowledge, this is the first study to measure molecular CMTs datasets to determine the association between higher stemness and basal-like tumors. Our findings are consistent with the results of Kim et al. (2020). They found that basal-like CMTs were enriched with the NMF3 metagene signature, which exhibited downregulation of claudin-encoding genes and E-cadherin (Kim et al. 2020), phenotypically similar to the low claudin breast cancer subtype, which is enriched with tumor-initiating cells and stem cells (Prat et al. 2010). However, in our study, we did not find an association between high stemness and the NMF3 metagene signature. Instead, the NMF2 subtype, considered a “DNA-repair” subtype, showed a higher mRNAsi and strong association with the stemness phenotype, opening new avenues for discussion and investigation. DNA damage and repair mechanisms play key roles in high stemness cells, such as stem cells and cancer stem cells. Both stem cells and cancer stem cells are endowed with effective DNA repair mechanisms to prevent the accumulation and propagation of genetic damage to daughter cells and to develop mechanisms of resistance to chemotherapy and radiotherapy, respectively (Vitale et al. 2017). This can explain the enrichment of the NMF2 subtype with high-stemness tumors in our study. In addition, a limitation of our study is that we could not understand the association between higher stemness, NMF metagene signatures, and histological types, since simple CMTs exhibiting higher stemness were more frequently classified in the NMF1 metagene signature (Kim et al. 2020). However, how stemness and the NMF metagene signatures correlate with therapy resistance in CMTs is open for future investigations.

Using a correlation analysis, we identified two possible new druggable targets for CMT. Both *POLA2* and *APEX1* were highly correlated with stemness. In addition, *POLA2* was highly expressed in simple CMTs than in other subtypes and normal samples, whereas *APEX1* expression was increased in general CMTs in comparison with normal samples. These translational and comparative analyses may contribute to drug repositioning and repurposing in CMTs, particularly targeting the stemness phenotype (Giuliano et al. 2023). In conclusion, exploiting the stemness phenotype in CMTs using omics and machine learning approaches holds promise for understanding the development of this heterogeneous disease and its subtypes, and for identifying novel therapeutic targets.

## Methods

### RNA expression data

RNA-seq data of canine mammary tumors were downloaded from the GEO database (GSE119810) (Kim et al. 2020). The database contains 157 tumor tissues and 64 matched, normal, and adjacent tissues. The original samples were classified according to grade, lymphatic invasion, ER status, HER2 score, Histology, NMF cluster, and PAM50 subtypes.

### Stemness index model

To predict the stemness phenotype in canine samples, we applied a machine learning algorithm described by Malta et al. (2018) to generate a one-class logistic regression (OCLR) algorithm employing human stem cell (iPS and ES) data from the Progenitor Cell Biology Consortium (PCBC) to estimate the stemness (mRNAsi) index in human tumors. The stem cell transcriptomic matrix used to train OCLR contained 12,945 gene expression values. Furthermore, this model was validated using leave-one-out cross-validation, which leaves one stem cell out of training, and then applies the model to non-SC cells (their differentiated ecto-, meso-, and endoderm progenitors). Moreover, Malta’s model presented a high correlation among other cancer stem cell signatures, providing significant insights into the biological and clinical features of pan-cancer. The workflow to generate the mRNAsi is available at https://github.com/ArtemSokolov/PanCanStem.

### Stemness index prediction

First, we converted the gene IDs from Canis lupus familiaris to Homo sapiens using tables of Ensembl data downloaded via the BioMart mining tool (https://www.ensembl.org/biomart/martview/). We then mapped ENSEMBL gene ID to gene symbols using the org.Hs.eg.db package from the Bioconductor repository. Genes with no such intersections were excluded. The previously described prediction stemness index model derived from pluripotent cells (Malta et al., 2018) was applied to the gene expression profiles of canine and human mammary cancer cell lines, leading to their mRNAsi. The indices were scaled to 0-1 (0 being a low stemness score and 1 being a high score) by subtracting the minimum and dividing by the maximum. The top 439 genes of the stemness signature (439 genes) were selected based on the Euclidean distance of the absolute weight of the stemness signature model. From the 12,953 genes in the original model, we applied 11,251 genes owing to the conversion gene IDs output. In relation to stemness prediction in human cell lines and canine stem cells, we applied 12,257 and 7,929 genes, respectively, as a result of the number of genes in common among RNA-seq from GEO datasets.

### Correspondence analysis

Simple (CA) and multiple (MCA) correspondence analyses were performed in the RStudio 4.3.1 environment using the packages FactoMineR (Lê et al., 2008) and cabootcrs (https://CRAN.R-project.org/package=cabootcrs). Contingency tables of the categorical variables were created, and to study the associations between variables, a chi-square test with k degrees of freedom, followed by the analysis of the adjusted standardized residuals, was performed. The Chi-squared test evaluates whether the associations between categorical variables are randomly associated. Adjusted standardized residuals higher than 1.96 indicate an association between variables in the matrix. To perform CA, the categorical variables should not be randomly associated. To create the perceptual map, inertia was determined as the total chi-squared divided by the number of samples, resulting in the number of associations in the dataset. MCA was performed based on the binary and Burt Matrices. Row and column profiles were determined to demonstrate the influence of each category of variables on the others. Matrices were defined based on the row and column profiles. Eigenvalues were then extracted to represent the number of dimensions that could be captured in the analysis. Finally, the x- and y-axis coordinates of the perceptual map were determined, allowing the category of the variables to be represented and established.

### Correlation analysis

To identify genes directly related to the Stemness Index obtained in carcinoma and normal samples, a linear correlation analysis was performed between the Stemness Index and the expression of each gene individually, using the cor.test function of the stats package. Spearman method was applied considering results with “p.value<0.05” and correlation “-0.5>correlation>0.5”. The 50 genes with the highest positive correlation and the 50 genes with the lowest negative correlation between the Stemness Index and the expression values are indicated in the heatmap.

### Statistical analysis

The stemness Index was compared between multiple groups using one-way ANOVA with post-hoc Tukey’s test. An unpaired T-test was used to compare the stemness index between the two groups. Correlation analysis was performed using the Spearman correlation coefficient. Differences were considered significant when the p < 0.05.

### Data access

All raw and processed RNA sequencing data used in this study was submitted to the NCBI Gene Expression Omnibus (GEO) under accession number GSE119810 (Kim et al., 2020).

## Acknowledgements

This study has been supported by grants from the Sao Paulo Research Foundation (FAPESP) (2018/00583-0, 2019/14928-1, 2021/00283-9, 2022/06305-7, 2023/05099-7, 2023/07358-0).

## Author contributions

PLPX, MM, RLSS, MEGJ, and TMM conducted the study, contributing with concept/design, acquisition of data, and data analysis/interpretation. PLPX, MM, RLSS, and MEGJ created images and figures. PLPX wrote the first version of the manuscript. HF, RFS and TMM contributed with interpretation of data, discussion of data, and corrected the final version of the manuscript.

